# Sale of critically endangered sharks in the United States

**DOI:** 10.1101/2023.07.30.551124

**Authors:** Savannah J. Ryburn, Tammy Yu, Kelly Jia-Wei Ong, Meggan A. Alston, Ella Howie, Peyton LeRoy, Sarah Elizabeth Giang, William Ball, Jewel Benton, Robert Calhoun, Isabella Favreau, Ana Gutierrez, Kayla Hallac, Dakota Hanson, Teagan Hibbard, Bryson Loflin, Joshua Lopez, Gracie Mock, Kailey Myers, Andrés Pinos-Sánchez, Alejandra Maria Suarez Garcia, Adriana Retamales Romero, Audrey Thomas, Rhiannon Williams, Anabel Zaldivar, John Francis Bruno

**Affiliations:** Environment, Ecology, and Energy Program, The University of North Carolina at Chapel Hill, Chapel Hill, North Carolina, 27599-3275 USA; The Department of Biology, The University of North Carolina at Chapel Hill, Chapel Hill, North Carolina, 27599-3280 USA; Colegio de Ciencias Biológicas y Ambientales, Universidad San Francisco de Quito, Diego de Robles y Pampite, 170901, Quito, Ecuador

**Keywords:** Shark meat, mislabeling, DNA barcoding, shark conservation, wildlife trade

## Abstract

Shark meat is widely available in the United States in grocery stores and seafood markets. The meat is often mislabeled or generically labeled as “shark”. The ambiguity of these generic labels makes it challenging to assess the conservation implications of this practice and for consumers to avoid species with high mercury concentrations. For this study we purchased and DNA barcoded 30 shark products purchased in the United States to determine their species identity and conservation status. These samples consisted of 19 filets sold in grocery stores, seafood markets, and Asian specialty markets (mostly in North Carolina) and 11 ordered online as “jerky”. 70% of samples were “soft mislabeled” (i.e., labeled generically as shark but not as a specific species). Of the nine samples labeled to species, eight were mislabeled (e.g., spinner shark labeled as mako shark). Only one sample was correctly labeled. All 30 samples were identified as shark and came from 11 different species, including three species listed by the IUCN as Critically Endangered: great hammerhead, scalloped hammerhead, and tope. The first two species have been found to contain very high levels of mercury, illustrating the implications of seafood mislabeling for human health. The widespread availability of shark meat in U.S. grocery stores is surprising given the dramatic decline of shark populations globally. Moreover, the fact that nearly all shark meat is either mislabeled or not labeled to species amplifies the problem. Accurate, verified product labels for shark meat would benefit consumers and shark conservation efforts, and should be a priority for the seafood industry.

## Introduction

Fishing has caused dramatic declines in shark populations worldwide (Baum et al., 2003; Ward-Paige et al., 2010; Valdivia, Cox, & Bruno, 2017; Roff et al., 2018). As a result, one-third of shark species are threatened with extinction and listed as “critically endangered”, “endangered”, or “vulnerable” by the International Union for Conservation of Nature (IUCN) (Dulvy et al., 2021). Sharks are generally harvested for food, and in many regions they are targeted specifically for their culturally and economically valuable dorsal fins (Clarke, 2004; Man, Wu, & Wong, 2014). Shark fins are typically served as soup throughout Asia. Consumers perceive shark fin as a luxury item associated with class, believed to have positive health benefits (Man, Wu, & Wong, 2014).

Most shark products, especially fins, are exported to countries in East and Southeast Asia such as China, Hong Kong, Taiwan, Singapore, Malaysia, and Vietnam (Vannuccini, 1999). However, shark meat is also sold in many other countries and regions, where it is often sold legally and typically incorrectly labeled or generically labeled as “shark” (i.e., “soft mislabeled”) (Fig. 1). Shark meat is also commonly sold under various ambiguous terms to obscure its identity from consumers (Bornatowski, Braga, & Vitule, 2013). For example, in South Africa shark is sold as “ocean filets’’ or “skomoro”, in Brazil elasmobranchs are sold as “cação” (Bornatowski, Braga, & Vitule, 2013), in Australia shark is sold under the term “flake” (Braccini et al., 2020), and in the United Kingdom small sharks are commonly sold as “rock salmon”, “huss”, “rock eel”, and “rigg” (Hobbs et al., 2019).

**Fig. 1.**
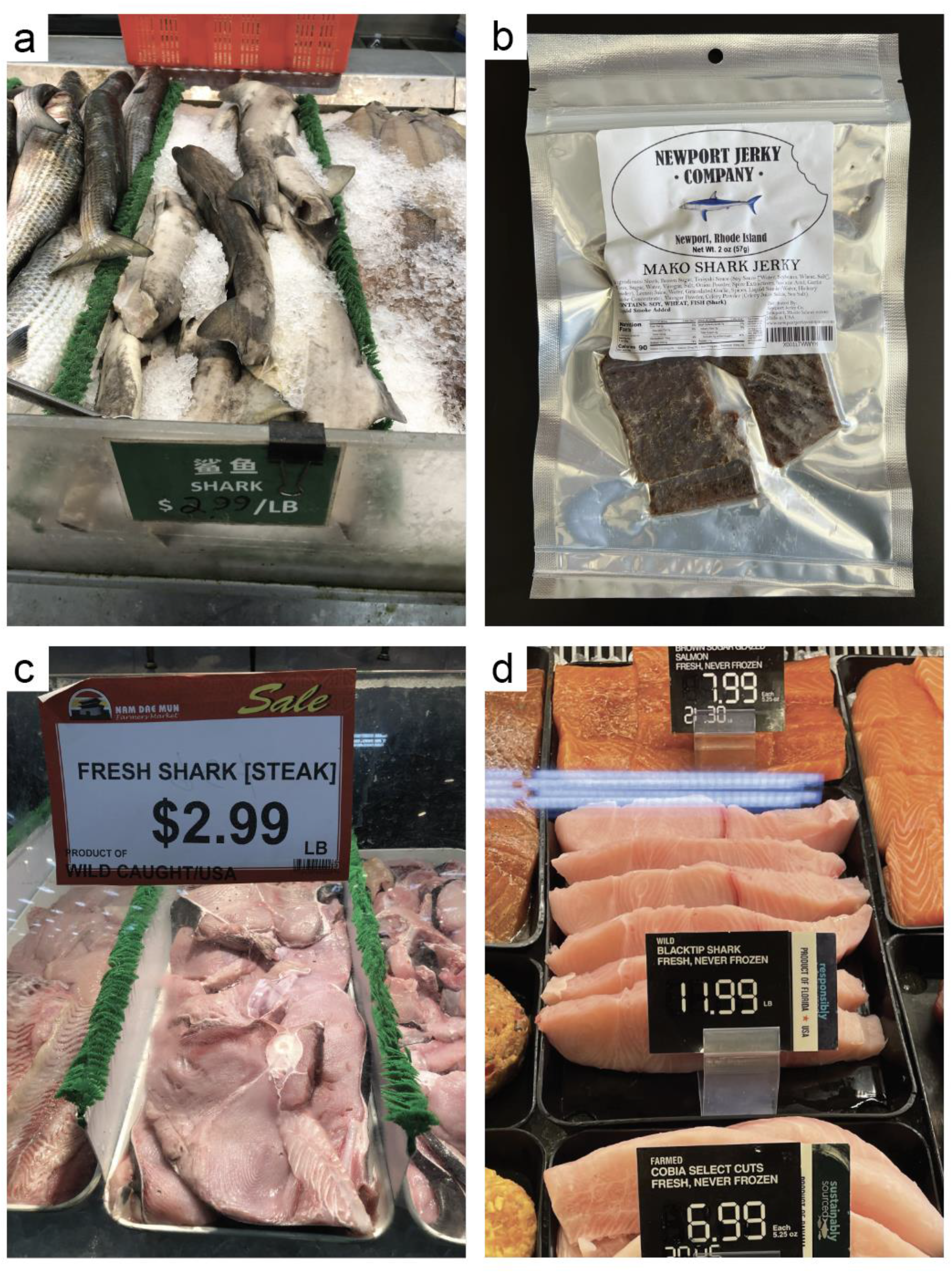
Photos exemplifying how some of the samples were labeled and displayed when purchased. (a) Dusky smooth-hound sold whole, missing the head, labeled as “shark” in English and Chinese at an Asian grocery store in Orlando, FL. (b) Shortfin mako shark sold as jerky and purchased online from the Newport Jerky Company in Rhode Island labeled as “mako shark jerky”. (b) Common thresher shark sold as “fresh shark [steak]” at an Asian grocery store in Duluth, GA. The label also included “wild caught/USA”. (d) Blacktip shark labeled as “wild blacktip shark, fresh never frozen” from a Publix grocery store in Cary, NC. The label also included that it was a “product of Florida” and that it was “responsibly sourced”.

Some of the world’s largest consumers of shark meat are actually South America and Europe, with the largest importers being Italy, Brazil, Uruguay, and Spain (Dent & Clarke, 2015). In Europe and North America, few countries restrict the sale of even species categorized as “critically endangered” by the IUCN. Consumers in these regions could regularly consume shark without even realizing it. For example, Bornatowski et al. (2015) interviewed customers at fish markets in Southern Brazil and found that 61% of respondents claimed that they ate cação but did not eat shark. Hobbs et al. (2019) reported that shark (primarily spiny dogfish *Squalus acanthias*) is commonly sold in takeaway fish and chips meals across the United Kingdom. They also reported that the sale of shark is common in UK grocery stores and seafood markets including IUCN critically endangered scalloped hammerhead (*Sphyrna lewini*). Likewise, Munguia-Vega et al. (2022) found that in Mexico, scalloped hammerhead and several other IUCN endangered and vulnerable species were being sold generically as “cazon” (a term used in the Mexican seafood trade to mean any shark meat). Generic umbrella terms allow for species that are subject to international trade restrictions under the Convention on International Trade in Endangered Species (CITES) to enter the market unnoticed.

Although shark appears to be widely traded in the global food system, very little information is available about the frequency, geographic extent, species identity, and numerous other aspects of the shark trade. Therefore, it is impossible to quantitatively assess the implications of this practice for either human health or marine biodiversity conservation. Hasan et al., 2023 reported that there are only 33 studies that have used molecular techniques to determine shark product mislabeling, and of these only three were conducted in the United States. The purpose of this study was to quantify the identity of “shark” sold in the United States. We purchased 30 samples from grocery stores and online jerky vendors. Using standard DNA barcoding techniques, we identified each sample to species. We found that critically endangered sharks, including great and scalloped hammerheads, are available in grocery stores. Moreover, of the 30 samples we collected, 97% were mislabeled.

## Methods

### Sample collection

This study was conducted by students and instructors (including undergraduate teaching assistants) in the Seafood Forensics course (BIOL 221) at The University of North Carolina at Chapel Hill. A total of 30 shark products were collected from September 2021 to September 2022, including 19 raw shark steaks and 11 packages of shark jerky from seafood markets, grocery stores, Asian markets, and online in the United States. All raw shark steaks were purchased on the east coast of the United States (i.e., North Carolina (*n* = 13), Florida (*n* = 3), Georgia (*n* = 2), and Washington DC (*n* = 1)). For each sample, we recorded how the seller had labeled the item (e.g., “shark steak”, “Mako shark”, “jerky”, etc.) and the price. Steak samples were kept frozen at -20 °C and jerky samples were kept at room temperature until DNA extraction.

### DNA Analysis

DNA extractions were completed using the Qiagen DNeasy Blood and Tissue Kit, following the manufacturer’s protocol but using a one hour digestion at 55°C and eluted with 25 μl of molecular water. Polymerase Chain Reaction (PCR) was used to amplify a fragment of the cytochrome c oxidase 1 (CO1) gene (Ivanova et al., 2007), cytochrome B (CytB) gene (Wolf et al., 2000), and NADH dehydrogenase subunit 2 (ND2) gene (Vella, Vella, & Schembri, 2017) for each sample (Table 1). We used all three fish/shark specific primers for each sample because, in some instances, the CO1 primer was not specific enough to determine the species. We added 20 - 100 ng of DNA per PCR reaction to separate 0.2 ml illustra puReTaq Ready-To-Go PCR bead tubes, along with 1.3 μl of each primer. To bring the overall volume to 25 μl, 18.8 μl of molecular grade water was added to the PCR bead tubes with the CO1 primer and 21.4 μl was added to the PCR bead tubes with the CytB and ND2 primers. Once the PCR bead was dissolved, we placed the tubes into a Bio-Rad T100 Thermal Cycler and used the protocol from (Spencer & Bruno, 2019).

**Table 1.**
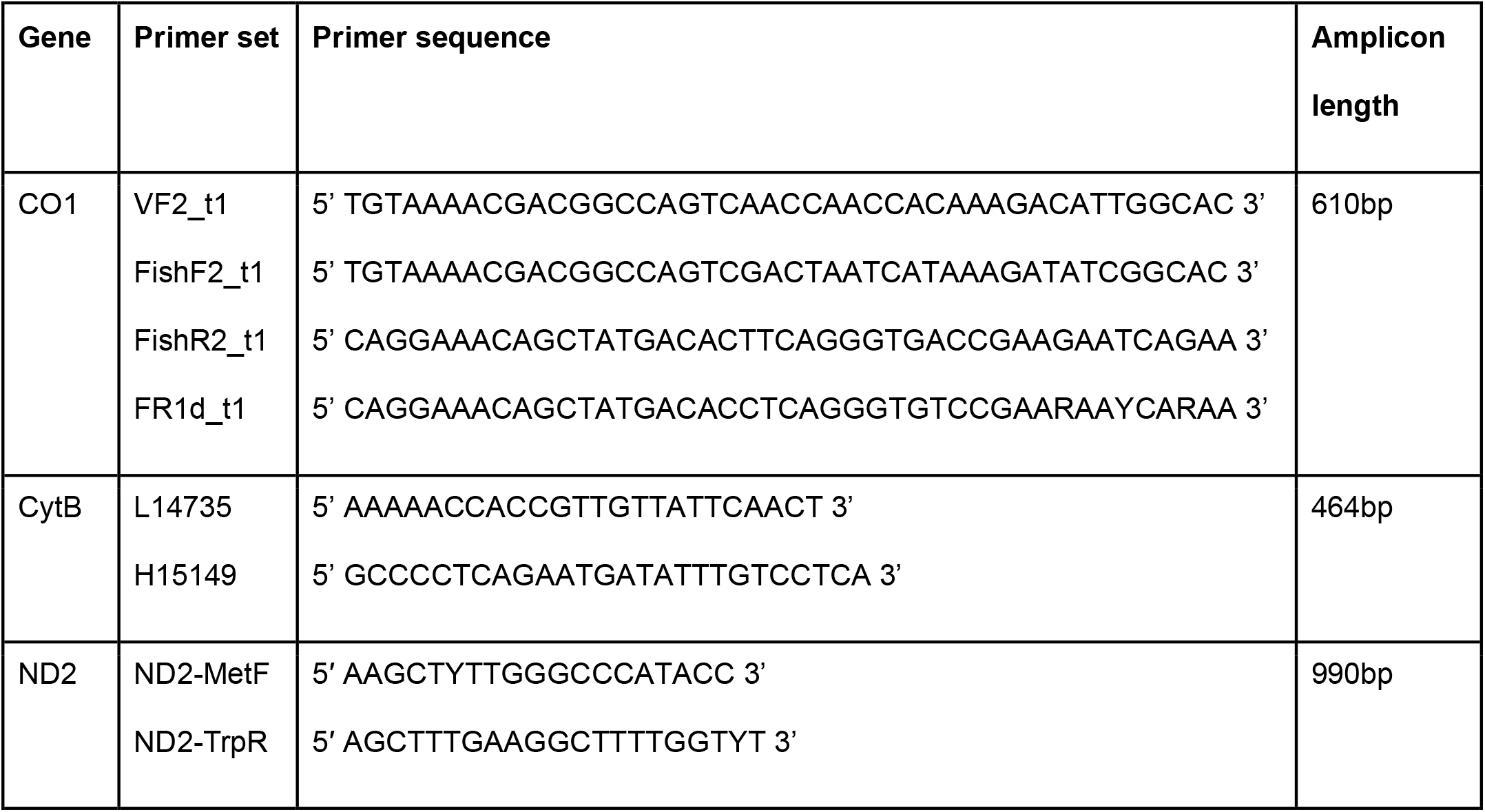
Primer sets used for species identification of shark samples.

### Sequencing results and analysis

The PCR products were purified, and Sanger sequenced in one direction by Eton Bioscience in Durham, North Carolina. The samples amplified with CO1 were sequenced using Eton’s universal primer M13F-21, the samples amplified with ND2 were sequenced with ND2-MetF, and the samples amplified with CytB were sequenced with L14735 (Table 1). To identify each sample to species level we used the Basic Local Alignment Search Tool (BLAST) on the National Center for Biotechnology Information (NCBI) website. We compared the results from the three primers to ensure that they matched and identities with a 94 percent identity or above were considered a positive identification. We chose the cut off of 94% (all but one jerky sample had a percent identity of 97% or higher) because about a third of our sample size was jerky that had been dehydrated and seasoned, meaning that we were working with degraded DNA. Each sample was extracted, PCRed, and sequenced twice to ensure accuracy.

Shark is frequently mislabeled in two ways. First, when it is labeled and sold as one species but is actually another species, oftentimes another species of shark (e.g., sold as blacktip shark but was actually lemon shark). We considered this “mislabeling” or “hard mislabeling”. It is also common for shark to be generically labeled as “shark steak” or “shark jerky” with no species specific labeling (Fig. 1). This is termed “soft mislabeling” and is common for other types of seafood as well (e.g., “scallops” or “squid”). Samples that were labeled as “mako shark” were also considered to be soft mislabeled because there are two different species of mako sharks (i.e., *Isurus oxyrinchus* and *Isurus paucus*). All samples were categorized as “soft mislabeled” (labeled as shark but not to species), mislabeled (labeled as a specific species but was not actually that species), or correctly labeled (labeled as a specific species and was actually that species). The conservation status of each species was determined using the IUCN Red List of Threatened Species (http://www.iucnredlist.org).

## Results

All 30 samples were successfully identified to species. 21 were soft mislabeled: 5 of the 11 jerky samples and 16 of the 19 filets. Of the 9 samples labeled to species, 8 were mislabeled and most were originally sold as mako shark. Only one sample was correctly labeled. All 30 samples were identified as shark and came from 11 species, including three species listed by the IUCN as Critically Endangered: great hammerhead, scalloped hammerhead, and tope (Fig. 2, Supplement 1). The average price in USD of the fresh shark meat was $6.30/lb ± 2.88 with prices ranging from $2.99/lb to $11.99/lb. The average price of the shark jerky was $5.08/oz ± 0.99.

**Fig. 2.**
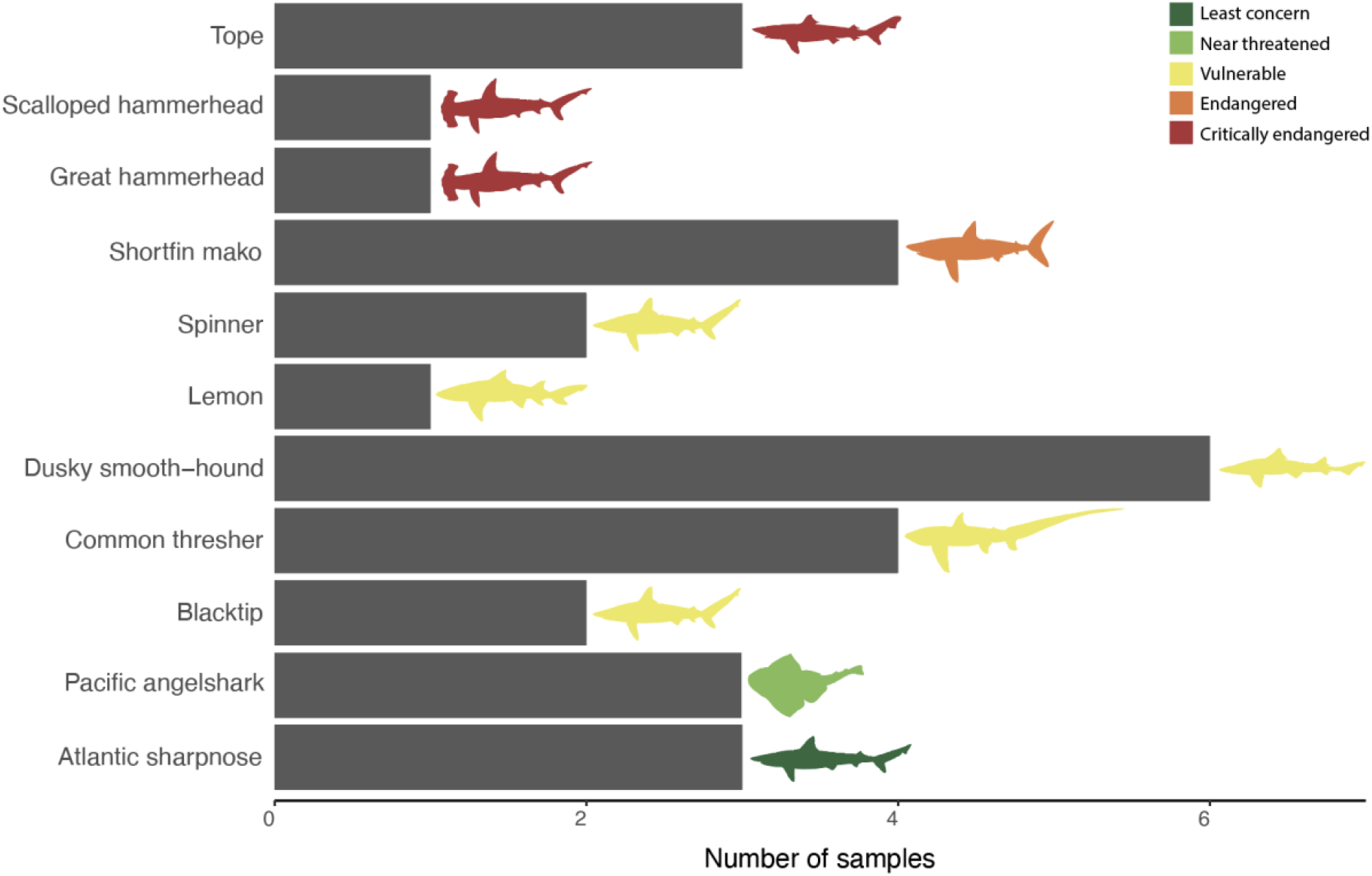
The number of each species found in our analysis using DNA barcoding to identify products labeled as shark. Species are color-coded to depict the IUCN status. Figure includes all samples (*n* = 30) and only the identity of each sample from the DNA barcoding results.

## Discussion

Overall, we found a high rate of shark product mislabeling and evidence of IUCN listed endangered species in the U.S. food supply. Thirty percent of our samples were classified by the IUCN as endangered or critically endangered species (Fig. 2). All 9 of these samples were also either mislabeled or soft mislabeled. Out of all 30 samples, only one was correctly labeled and it was from a blacktip shark (*Carcharias limbatus*). Although our sample size is relatively small (*n* = 30), we still detected multiple critically endangered and CITES Appendix II species being sold generically as “shark” in U.S. markets. These results resembled those of similar studies that occurred in Italy and Brazil. When using our study’s definition of soft mislabeling all 46 samples from Barbuto et al. (2010) in Italy would be considered soft mislabeled, while in Brazil, Almerón-Souza et al. (2018) collected 63 samples and 61 of them would be considered soft mislabeled and 2 were mislabeled.

Mislabeling and soft mislabeling can greatly impact the conservation of marine predators and the sustainability of the fishing industry. Although consumers are indeed being sold “shark”, there are more than 536 species of sharks (Dulvy et al., 2021) ranging in size, diet, habitat, and endangerment status. For example, scalloped hammerhead sharks are critically endangered with a population that continues to decline globally (Rigby et al., 2019). Several aspects of the life history of hammerhead sharks make them especially vulnerable to fishing (Gallagher et al., 2014). Despite their status, the scalloped hammerhead and other endangered top marine predators are frequently targeted, caught, and shipped internationally.

Shark fins are a highly sought out product throughout Asia (Eriksson & Clarke, 2015) specifically for shark fin soup. The lower caudal fin of hammerheads (*Sphyrna spp*.) and the shortfin mako (*Isurus oxyrinchus*) are considered to have the best quality fin needles for consumption (Clarke, Milner-Gulland, & Bjørndal, 2007), resulting in a high demand for these species. Cardeñosa et al. (2022) found that scalloped hammerhead was the third most common species sold in a Hong Kong fish market as fins. The shortfin mako was the sixth most common species (Cardeñosa et al., 2022). Additionally, Almerón-Souza et al. (2018) found scalloped hammerhead to be the most abundant species when testing shark meat sold in Brazil. These and other shark species threatened with extinction are present-to-common in the food supply in numerous countries around the world.

Eight of our 11 samples labeled to species were labeled as mako shark. A higher demand for mako shark fins and meat could be why most mislabeled meat products were labeled as mako. Makos (i.e., *Isurus oxyrinchus* and *Isurus paucus*), threshers (i.e., *Alopias spp*.), larger hammerheads (i.e., *Sphyrna lewini, Sphyrna mokarran*, and *Sphyrna zygaena*), and all requiem sharks (e.g., *Carcharhinus brevipinna, Negaprion brevirostris, Carcharhinus limbatus*, and *Rhizoprionodon terraenovae*) are all classified under CITES Appendix II which require export permits due to their extreme vulnerability if trade is not closely controlled. 60% of our samples were listed under CITES Appendix II. Mislabeling can be a tactic used to disguise endangered species (e.g., *Sphyrna lewini, Sphyrna mokarran*, and *Isurus oxyrinchus*) being harvested for the fin trade since different species of shark meat can look identical when fileted or dried as jerky.

The Shark Conservation Act of 2010 (H.R. 81. Shark conservation act of 2010, Public Law 111-348, 2010) requires that all sharks in the United States, with one exception (*Mustelus canis*), be brought to shore with their fins naturally attached. This law has forced fishermen to land the entire shark rather than only taking the fins and disposing the body at sea, thereby potentially increasing the amount of shark meat being sold as byproduct. Shark meat is oftentimes considered to be “junk meat” due to its unpleasant smell and taste which is a result of high amounts of urea (Suryaningsih et al., 2020). The market cost for most of our samples was very low, especially in grocery stores. For example, a scalloped hammerhead filet was only $4.99 / lb. Given the massive ecological cost of harvesting such an important, rare, and long-lived predator (e.g., scalloped hammerhead) it is remarkable how low their market price is.

Shark products can contain heavy metals including arsenic, mercury, and methylmercury (Garcia Barcia et al., 2020) with the highest concentrations in muscle tissue (Tiktak et al., 2020). Consumption of these heavy metals can have negative effects on human health (Tiktak et al., 2020; Souza-Araujo et al., 2021). For example, Garcia Barcia et al. (2020) found high levels of mercury in meat and fin products from large hammerhead species, *Sphyrna mokarran, S. zygaena*, and *S. lewini*, and advised consumers to avoid these species. They also recommended that “species specific advisories to be issued for meat and fin products from oceanic whitetip and dusky smooth-hound sharks, which should be avoided by women of childbearing age again due to high mercury levels” (García Barcia et al., 2022). We were sold three of these five species (i.e., scalloped hammerhead, great hammerhead, and dusky smooth-hound) all which were labeled generically as “shark”. Consumption of mercury and methylmercury can cause damage to the brain and central nervous system while arsenic can lead to skin, bladder, and lung cancer (Ratcliffe, Swanson, & Fischer, 1996; Ravenscroft, Brammer, & Richards, 2011; Skalny et al., 2022; Honda, Hylander, & Sakamoto, 2006). Consumption of all three of these metals has been related to fetal cognitive development complications and infant death (Farzan, Karagas, & Chen, 2013; Tolins, Ruchirawat, & Landrigan, 2014; Quansah et al., 2015; Honda, Hylander, & Sakamoto, 2006). The soft mislabeling of shark meat can be detrimental to human health. When consumers are purchasing mislabeled or soft mislabeled shark meat, they have no way to know what species they are consuming and what the associated health risks might be.

Our sampling and inferences about shark steak mislabeling are biased towards North Carolina. However, the jerky samples were purchased from national vendors (all based in the United States) therefore the inferences made from those samples are representative of the geographic region. Due to our limited sample size, future work should include a larger, more geographically widespread sampling to better characterize the sale of shark product. The species composition of shark meat being sold in the United States and the rest of the world is still largely unknown. Increased monitoring and accurate, verified product labels for shark meat are necessary given the high proportion of species threatened with extinction found within our small sample size. A relatively new technique known as genetic stock identification (Fields et al., 2020) could also be used to decipher what specific populations or regions the shark meat is coming from (e.g., scalloped hammerheads from the Eastern Pacific). This method could provide a better understanding of whether these CITES listed species are being imported illegally and if these sharks are being fished from overexploited regions.

## Acknowledgements

This study was funded in part by the Department of Biology at The University of North Carolina at Chapel Hill. Samples were collected and analyzed as part of the Seafood Forensics CURE class at UNC-CH (BIOL 221). The project was also partially funded by the National Science Foundation (OCE #2128592 to JFB).

## Author Contributions

*SJR, TY, KJO, MAA, EH, PL, SEG*, and *JFB* designed the study. *SJR, JFB, EH, PL, SEG, KJO, AT, GM, IJ*, and *AR* collected the samples. *SJR, TY, KJO, MAA, EH, PL, SEG, WB, JB, RC, IF, AG, KH, DH, TH, BL, JL, GM, KM, AP, AMS, AR, AT, RW*, and *AZ* completed the DNA analysis. *SJR* analyzed the data. The manuscript was written by *SJR* with help from *JFB. SJR, JFB*, and *MAA* revised the manuscript. All authors gave final approval for publication.

## Notes

### Competing Interest Statement

The authors have declared no competing interest.

