## Supplemental Table 1 for "Sale of critically endangered sharks in the United States"

**Supplement**

**Table S1.** List of what each sample was sold as, if the sample was mislabeled, soft mislabeled, or correctly labeled, and its conservation status. Table includes all samples (*n* = 30). * Signifies that the sample was ordered online.

| **Sample ID** | **Sold as** | **Mislabeling status** | **True Identity** | **Common name** | **IUCN status** | **Price** | **Location Purchased** |
|---|---|---|---|---|---|---|---|
| 1 | Shark steak | Soft mislabeled | *Carcharhinus brevipinna* | Spinner shark | Vulnerable | $5.99/lb | Cary, NC |
| 2 | Shark steak | Soft mislabeled | *Carcharhinus limbatus* | Blacktip shark | Vulnerable | $5.99/lb | Cary, NC |
| 3 | Shark loin | Soft mislabeled | *Negaprion brevirostris* | Lemon shark | Vulnerable | $8.05/lb | Raleigh, NC |
| 4 | Mako shark | Mislabeled | *Carcharhinus brevipinna* | Spinner shark | Vulnerable | $11.99/lb | Carrboro, NC |
| 5 | Mako shark | Soft mislabeled | *Isurus oxyrinchus* | Shortfin mako shark | Endangered | $11.99/lb | Knightdale, NC |
| 6 | Shark | Soft mislabeled | *Mustelus canis* | Dusky smooth-hound | Near threatened | $4.99/lb | Raleigh, NC |
| 7 | Blacktip | Mislabeled | *Isurus* | Shortfin | Endangered | $7.95/lb | Columbia, |

| | | | | | | | |
|---|---|---|---|---|---|---|---|
| | shark | | *oxyrinchus* | mako shark | | | NC |
| 8 | Shark | Soft mislabeled | *Mustelus canis* | Dusky smooth-hound | Near threatened | $4.99/lb | Raleigh, NC |
| 9 | Shark | Soft mislabeled | *Rhisoprionodon terranovae* | Atlantic sharpnose shark | Least concern | $2.99/lb | Raleigh, NC |
| 10 | Wild blacktip shark | Correctly labeled | *Carcharhinus limbatus* | Blacktip shark | Vulnerable | $11.95/lb | Cary, NC |
| 11 | Shark | Soft mislabeled | *Mustelus canis* | Dusky smooth-hound | Near threatened | $2.99/lb | Orlando, FL |
| 12 | Shark steak | Soft mislabeled | *Rhizoprionodon terraenovae* | Atlantic sharpnose shark | Least concern | $4.99/lb | Orlando, FL |
| 13 | No label | Soft mislabeled | *Rhizoprionodon terraenovae* | Atlantic sharpnose shark | Least concern | $3.99/lb | Orlando, FL |
| 14 | Fresh shark (steak) | Soft mislabeled | *Alopias vulpinus* | Common thresher shark | Vulnerable | $2.99/lb | Duluth, GA |

| | | | | | | | |
|---|---|---|---|---|---|---|---|
| 15 | Shark steak | Soft mislabeled | *Mustelus canis* | Dusky smooth-hound | Near threatened | $7.78/lb | Raleigh, NC |
| 16 | Shark steak | Soft mislabeled | *Sphyrna mokarran* | Great hammerhead shark | Critically endangered | $5.99/lb | Johns Creek, GA |
| 17 | Shark steak | Soft mislabeled | *Sphyrna lewini* | Scalloped hammerhead shark | Critically endangered | $4.99/lb | Charlotte, NC |
| 18 | Thresher shark jerky | Mislabeled | *Galeorhinus galeus* | Tope | Critically endangered | $16.99/2 oz | Newport, RI* |
| 19 | Shark steak | Soft mislabeled | *Mustelus canis* | Dusky smooth-hound | Near threatened | $4.99/lb | Greensboro, NC |
| 20 | Mako shark jerky | Soft mislabeled | *Alopias vulpinus* | Common thresher shark | Vulnerable | $14.99/3 oz | Carson City, NV* |
| 21 | Mako shark jerky | Mislabeled | *Isurus oxyrinchus* | Shortfin mako shark | Endangered | $14.99/3 oz | Newport, RI* |
| 22 | Mako shark jerky | Mislabeled | *Isurus oxyrinchus* | Shortfin mako shark | Endangered | $14.99/3 oz | Evansville, IN* |

| | | | | | | | |
|---|---|---|---|---|---|---|---|
| 23 | Sriracha shark jerky | Soft mislabeled | *Alopias vulpinus* | Common thresher shark | Vulnerable | $14.99/3 oz | Evansville, IN* |
| 24 | Mako shark jerky | Soft mislabeled | *Galeorhinus galeus* | Tope | Critically endangered | $17.99/3 oz | Evansville, IN* |
| 25 | Peppered shark jerky | Mislabeled | *Squatina californica* | Pacific angelshark | Near threatened | $16/2.75 oz | Fresno, CA* |
| 26 | Peppered shark jerky | Soft mislabeled | *Squatina californica* | Pacific angelshark | Near threatened | $14.95/2.75oz | Big Bear City, CA* |
| 27 | Mako shark | Mislabeled | *Galeorhinus galeus* | Tope | Critically endangered | $16.25/3 oz | Evansville, IN* |
| 28 | Shark steak | Soft mislabeled | *Mustelus Canis* | Dusky smooth-hound | Near threatened | $3.99/lb | Washington, DC |
| 29 | Shortfin mako shark jerky | Mislabeled | *Alopias Vulpinus* | Common thresher shark | Vulnerable | $17.99/2.75oz | Gulf Shores, AL* |
| 30 | Shark jerky | Soft mislabeled | *Squatina californica* | Pacific angelshark | Near threatened | $16.99/2.75oz | Myrtle Beach, SC* |
